# P95-HER2 promotes metastatic progression by biasing MRTFA dependent signaling

**DOI:** 10.64898/2026.07.22.739975

**Authors:** Joseph D. Fernandes, Erin Bresnahan, Hillary Zawada, Joshua D. Ginzel, H. Kim Lyerly, Bruce W. Rogers, Shyam M. Kavuri, Allison J. Introne, Melanie R. Sadecki, Hidetoshi Mori, Glenn Doherty, Alexander D. Borowsky, Kevin B. Flores, Jose Javier Bravo-Cordero, Joshua C. Snyder

**Affiliations:** Department of Cell Biology, Duke University Medical Center, Durham, NC, USA; Division of Hematology and Medical Oncology, Department of Medicine, The Tisch Cancer Center, Icahn School of Medicine at Mount Sinai, New York, NY, USA; Department of Surgery, Division of Surgical Sciences, Duke University Medical Center, Durham NC, USA; Department of Pathology, Duke University Medical School, Durham, NC, USA; Department of Immunology, Duke University School of Medicine, Durham, NC, USA; Lester Sue and Smith Breast Center, Department of Medicine, Baylor College of Medicine, Houston, TX, USA; Department of Mathematics, North Carolina State University, Raleigh, NC, USA; Department of Pathology, University of California San Francisco, San Francisco, CA, USA; Department of Pharmacology and Cancer Biology, Duke University School of Medicine, Durham, NC, USA; Microscopy and Advanced Bioimaging Core, Icahn School of Medicine at Mount Sinai, New York, New York, USA

**Author notes:** Correspondence to Jose Javier Bravo-Cordero PhD, Icahn School of Medicine at Mount Sinai, 1468 Madison Avenue, Box 1079, NY 10029 (Bravo-Cordero) and Joshua C. Snyder, Ph.D., Duke University Medical Center, Box 2606, Durham, NC 27710. Equal Contributions (Joseph D. Fernandes, Erin Bresnahan).

## Abstract

Naturally occurring isoforms of the proto-oncogene Erb-B2 Receptor Tyrosine Kinase 2 (ERBB2, HER2) incite unique pathways of tumor progression in mouse models of breast cancer. Although each isoform has been shown to progress to metastasis, the N-terminally truncated isoform (p95) was previously shown to be biased toward early distant dissemination prior to detection. Here we show how HER2 isoforms differentially promote go-or-grow phenotypes through biased utilization of tyrosine autophosphorylation sites in the intracellular tail of HER2. Fluorescently barcoded humanized full-length HER2 (WT), exon-16 splice HER2 isoform (d16), and p95 expressing tumor cell lines were derived from HER2 Crainbow mice, then utilized for a series of functional assays including proliferation, tumor growth rate, cell motility, collective cell migration, cellular morphology, and invasive potential. Quantitative analysis reveals biased tumor cell behaviors across each genotype. WT tumor cells are biased toward proliferation and collective migration, d16 cells are biased toward proliferation and individual motility, and p95 tumor cells are biased toward individual motility and invasion. Single cell analysis reveals a myogenic-like state transition in p95 cells accompanied by an increased nuclear translocation of the Myocardin Related Transcription Factor A (MRTFA). MRTFA knockdown, as well as knockdown of a downstream effector, Transforming growth factor beta 1 induced transcript 1 (TGFB1I1), both inhibit the p95 invasive phenotype. Furthermore, intracellular residue tyrosine 1139 (Y1139) is necessary for MRTFA translocation, and when edited in p95 cells (Y1139F) invasion and motility phenotypes are subsequently lost. Our data illustrate the importance of functionally selective signaling in HER2 and highlight the need for therapeutics that intercept metastasis by targeting HER2 biased signaling.

## Introduction

Early dissemination of breast cancer cells before screen detection is a major contributor to metastasis and poor prognosis (1–6). HER2+ breast cancer cells are particularly prone to early spread, yet the receptor-level signaling mechanisms underlying this phenomenon are not yet fully understood (7,8). Recent evidence suggests that naturally occurring HER2 isoforms may differentially promote invasion and motility (‘go’ behaviors) or proliferative signaling (‘grow’ behaviors) (9,10). Here we compare wild-type (WT) HER2, an exon 16 null isoform of HER2 (d16), and N-terminally truncated HER2 (p95) to determine how isoform-specific receptor signaling promotes functionally selective signaling and hallmark cancer behaviors like “sustaining proliferative signaling” and “activating invasion and metastasis” (11).

HER2 is a receptor tyrosine kinase (RTK) lacking an orthotopic ligand and signals by forming heterodimers with ligand-bound HER3. This results in HER2-driven tyrosine kinase activity, transautophosphorylation of tyrosine residues in the carboxyl terminal tail, adaptor recruitment (i.e. GRB2, SHC), and downstream signaling (12,13). In contrast to WT HER2, d16 and p95 isoforms can homodimerize and signal independently of HER3 to exert potent tumorigenic potential (14–18). d16 HER2 is an alternative splicing product lacking exon 16, which encodes a 16 amino acid N-terminal/juxtamembrane domain, that is expressed in multiple human cancers, including breast cancer. Murine models have demonstrated that d16 HER2 is associated with luminal phenotypes that are sensitive to HER2-targeted monoclonal antibody therapy but also can progress to metastasis (16,17,19–21). p95 HER2 is formed due to either alternative translation at an internal start codon (amino acid 611) or due to ADAM10-mediated cleavage of a nearby protease cleavage site. Either mechanism generates an N-terminal truncation devoid of the trastuzumab binding epitope, rendering this isoform resistant to HER2-targeted monoclonal antibody therapy (22–24). HER2 has also been shown to drive tumor cell mediated inflammatory responses (25). Similarly, p95 may also drive paracrine activation of additional signaling pathways, like JAK/STAT signaling, and ultimately immune senescence that can also inhibit HER2-targeted antibody drug conjugates (26) leading to treatment resistant metastatic breast cancer (27–33).

Previously we described a Cancer rainbow model to fluorescently barcode and directly compare HER2 isoforms in the same mouse to better understand how each isoform impacts breast cancer progression (9,10,34). In this HER2 Crainbow model system, WT HER2 tumors are rare compared to the extensive d16 or p95 tumors that frequently progress to metastasis. Surprisingly however, d16 and p95 tumors differ in their invasive potential early in malignant progression, with p95 exhibiting early invasion and dissemination prior to palpation whereas early d16 tumors are non-invasive and less likely to disseminate prior to detection (9,10).

Our goal here is to determine how isoforms of HER2 could differentially bias proliferation and invasion behaviors during the progression to treatment resistant breast cancer. Functional selectivity is a phenomenon most often studied in the context of the G protein-coupled receptor superfamily, where the same receptor can engage distinct downstream effectors and bias signaling pathways (35). Functionally selective signaling is an emerging paradigm in RTKs (36,37), and in the case of HER2, unique transphosphorylation in the Carboxyl-tail may control direct or indirect engagement of multiple effectors (GRB2/SHC) to bias proliferative and invasive tumor cell behaviors (13,38–41). These studies led us to hypothesize that the d16 and p95 isoforms may exert distinct dissemination programs, with increased invasive mechanisms driven by p95 HER2. Herein we trace cell motility and invasion programs using cell models derived from HER2 Crainbow mice to elucidate biased signaling properties of HER2 isoforms. We show how each isoform drives unique tumor cell behaviors and specifically demonstrate how p95 HER2 differentially biases MRTFA/TGFB1I1 pathway through Y1139 in the C-terminal tail of HER2 to promote invasive behavior.

## Results

### Derivation of HER2 Crainbow cell lines

In order to investigate how specific HER2 isoforms regulate tumor cell phenotypes, we first developed primary cell lines derived from HER2 Crainbow (HER2BOW) tumors that express fluorescently barcoded WT-HER2 (mTFP1:Cyan), d16 HER2 (EYFP:yellow), and p95-HER2 (mKO:magenta) isoforms (9,10). HER2BOW tumors were mechanically and enzymatically digested, then culture purified to establish WT-HER2 (mTFP1:Cyan), d16 HER2 (EYFP:yellow), and p95-HER2 (mKO:magenta) expressing cell lines, herein referred to as HCR-WT (<u>H</u>ER2 <u>C</u>rainbow - <u>WT</u>), HCR-16 (<u>H</u>ER2 <u>C</u>rainbow -d<u>16</u>), and HCR-95 (<u>H</u>ER2 <u>C</u>rainbow-p<u>95</u>) (Figure 1A). Each isoform-specific cell line maintained fluorescent protein expression and expression of human HER2 as confirmed by both N-terminus and C-terminus immunostaining (Figure 1B). The HCR-95 cell line showed only C-terminal HER2 staining, as expected with the absence of its extracellular N-terminal domain. Cell lines were able to grow *in vitro* at similar rates of proliferation and population doubling (Figure 1C, D); however, HCR-WT cells were more densely-packed as quantified by an overall higher carrying capacity and exhibited a cobblestone phenotype typical of epithelial cells, as compared to both HCR-16 and HCR-95 which displayed larger, elongated cells (Figure 1B, E). To test growth behavior *in vivo*, cell lines were transplanted orthotopically into the mammary fat pad of NOD/SCID mice (Figure 1F). Each HER2BOW cell line was able to develop palpable tumors within 15 days at 100% penetrance (Figure 1G).

**Figure 1.**
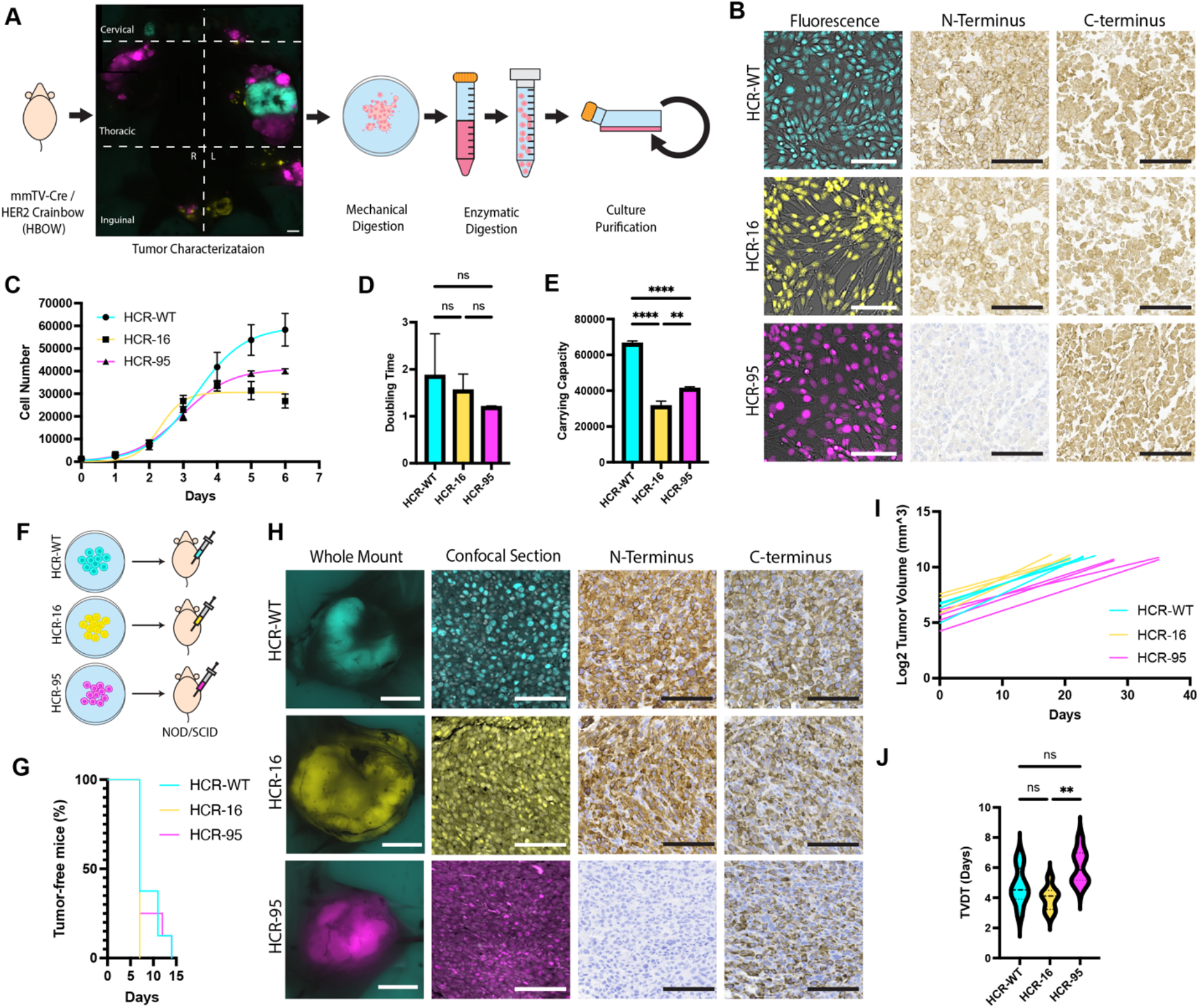
Development and characterization of HBOW cell lines. **A,** Schematic depicting isolation and purification process of HBOW cell lines. **B,** Confocal imaging of HCR cell lines in 2D culture (image shows overlay of fluorescence and DIC). IHC staining for N-terminus and C-terminus domains of HER2 receptor on HCR cell pellets. **C,** Cell proliferation *in vitro* as quantified by daily imaging and cell segmentation. Three independent experiments were averaged and fit with a logistic growth model. **D,** Population doubling time and **E,** Carrying capacity calculated from “C”. **F,** Schematic depicting orthotopic transplant of cells into NOD/SCID mice. **G,** Tumor-free survival of transplants by genotype (n=10 for each genotype). **H,** Fluorescent microscopy of isolated transplanted tumors. Whole mount tumor imaging (epifluorescent dissecting microscope) and tissue section imaging (confocal microscopy) are shown. IHC staining of N-terminus and C-terminus domains of HER2 on FFPE sections of transplant tumors. **I,** Linear curve fit of log2 transformed overall tumor volume. **J,** Tumor volume doubling time (TVDT) from “I”. Statistical significance was determined by one-way ANOVA with Tukey’s multiple comparisons test. ns, not significant; *p<0.05; **p<0.01; ***p<0.001; ****p<0.0001. Error bars are represented as mean±s.e.m. Scale bars: 5000 *μm* (A, H fluorescence-whole tissue), 100*μm* (B, H fluorescence-section, N-terminus, C-terminus), (mTFP1:cyan:WT HER2, EYFP:yellow:d16 HER2, and mKO:magenta:p95 HER2)

Furthermore, as observed *in vitro*, each tumor retained its fluorescent protein expression and predicted N-terminal and C-terminal HER2 immunostaining (Figure 1H). Although tumor penetrance was similar, tumor growth rates were noticeably different between the genotypes. Tumor growth was slower for HCR-95 transplants compared to HCR-WT and HCR-16 (Figure 1I). Furthermore, tumor volume doubling time for HCR-95 tumors was significantly higher than HCR-16 and trended higher than HCR-WT (P = 0.0660) (Figure 1J). Taken together, these data suggested an increased proliferative (“grow”) behavior of HCR-WT and HCR-16 cells as compared to HCR-95 cells.

### Differential Motility and Invasion Potential of HER2 isoforms

We next aimed to more quantitatively visualize motility differences, as part of a “go” phenotype, between the isoforms in an *in vitro* system. We first assessed kinematic motility by imaging live cell cultures for each cell line in a mixed 2-dimensional culture over the course of 16 hours, across three different cell culture substrates: normal tissue culture plastic, Fibronectin, and poly-D-lysine. To quantify motility behaviors, we employed a segmentation and tracking pipeline using both automated and generative artificial intelligence (AI) approaches (Figure 2A, Supplementary Methods). After imaging, generative AI was used to interpolate cell tracks. Following this, Cellpose was used to segment cells from each genotype, then nuclei were tracked using Trackpy. This resulted in all cells in a given image being tracked individually with great accuracy (Figure 2 B, C, Supplemental Figure S6, S8). Single cell tracks were used to calculate three population level motility metrics: average speed, persistence, and mean square displacement (MSD) slope. We also calculated two spatial metrics in order to investigate how cell genotypes in a neighborhood of 50µm interacted: this included a within-genotype correlation (i.e. “Self-correlation”, Figure 2D, Supplementary Methods), and cells without a neighbor (i.e. “Zero Neighbors”, Figure 2D, Supplementary Methods).

**Figure 2.**
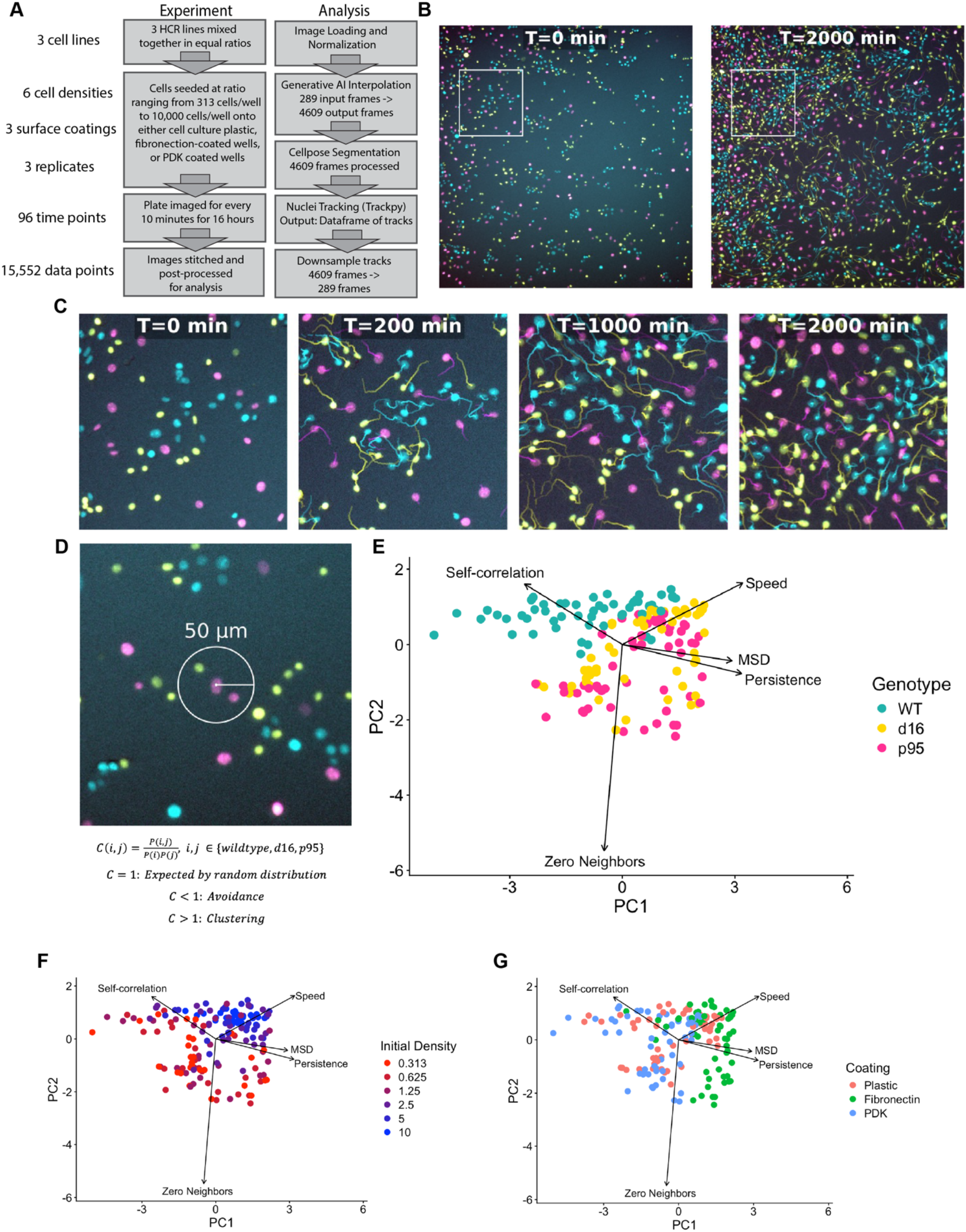
Timelapse live imaging and single cell tracking for motility analysis of HER2 Crainbow cell lines. **A,** Schematic of experimental conditions and cell tracking approach. **B,** Snapshots of a representative well seeded on tissue culture plastic at 10,000 initial cell density with single cell tracks overlaid. **C,** Region of interest from “B”. **D,** Diagram of spatial correlation analysis using a 50µm neighborhood. **E,** Principal component analysis (PCA) of motility and spatial crowding metrics across all conditions and colored by genotype. **F&G,** PCA of motility and spatial crowding metrics across all experiments colored by initial cell seeding density (F) and surface coating (G).

Principal Component Analysis (PCA) was performed on the population level motility (average speed, MSD, Persistence, and spatial interactions) to summarize variation across genotypes and experimental conditions (Figure 2E-G). HCR-16 and HCR-95 cells cluster similarly and are driven by their similarity in Speed, MSD, Persistence, and individual behavior (low self-correlation and zero neighbors), whereas HCR-WT is distinct for its collective cell behavior (high self-correlation) (Figure 2E). Moreover, HCR-16 and HCR-95 appeared most divergent from HCR-WT at lower seeding densities (Figure 2F). Although these findings were similar across three culture substrates, Fibronectin was associated with increased MSD and Persistence (Figure 2G). We performed additional statistical analysis using mixed-effects modeling to investigate significant differences across each genotype and condition which are reported in Supplemental Tables S1-S5. Altogether, the pairwise analysis again suggests that HCR-16 and HCR-95 share an increased persistence and diffusive potential compared to HCR-WT cell lines.

Next, we compared cell morphology characteristics indicative of invasive phenotypes. To visualize cell morphology, we stained each cell line with phalloidin and discovered HCR-95 cells exhibited a strikingly different actin cytoskeletal organization. HCR-95 cells were significantly larger than HCR-WT and HCR-16 cells, with a more spindle-like, elongated shape (Figure 3 A, B). HCR-95 cells also displayed thick aligned actin stress fibers that were significantly longer than that of HCR-WT and HCR-16 cells (Figure 3C). We also observed invadopodia-like actin-rich protrusions which have been shown to drive matrix degradation in cancer cells, identified as cortactin/F-actin positive puncta overlying points of focal matrix degradation (Figure 3D) (42–44), and found that HCR-95 cells show significantly more cortactin/F-actin positive puncta (Figure 3E, F). HCR-95 cells also developed large aggregates of up to hundreds of invadopodia organized into superstructures which appeared similar to previously described podosome/invadopodia rosettes (Figure 3G).

**Figure 3:**
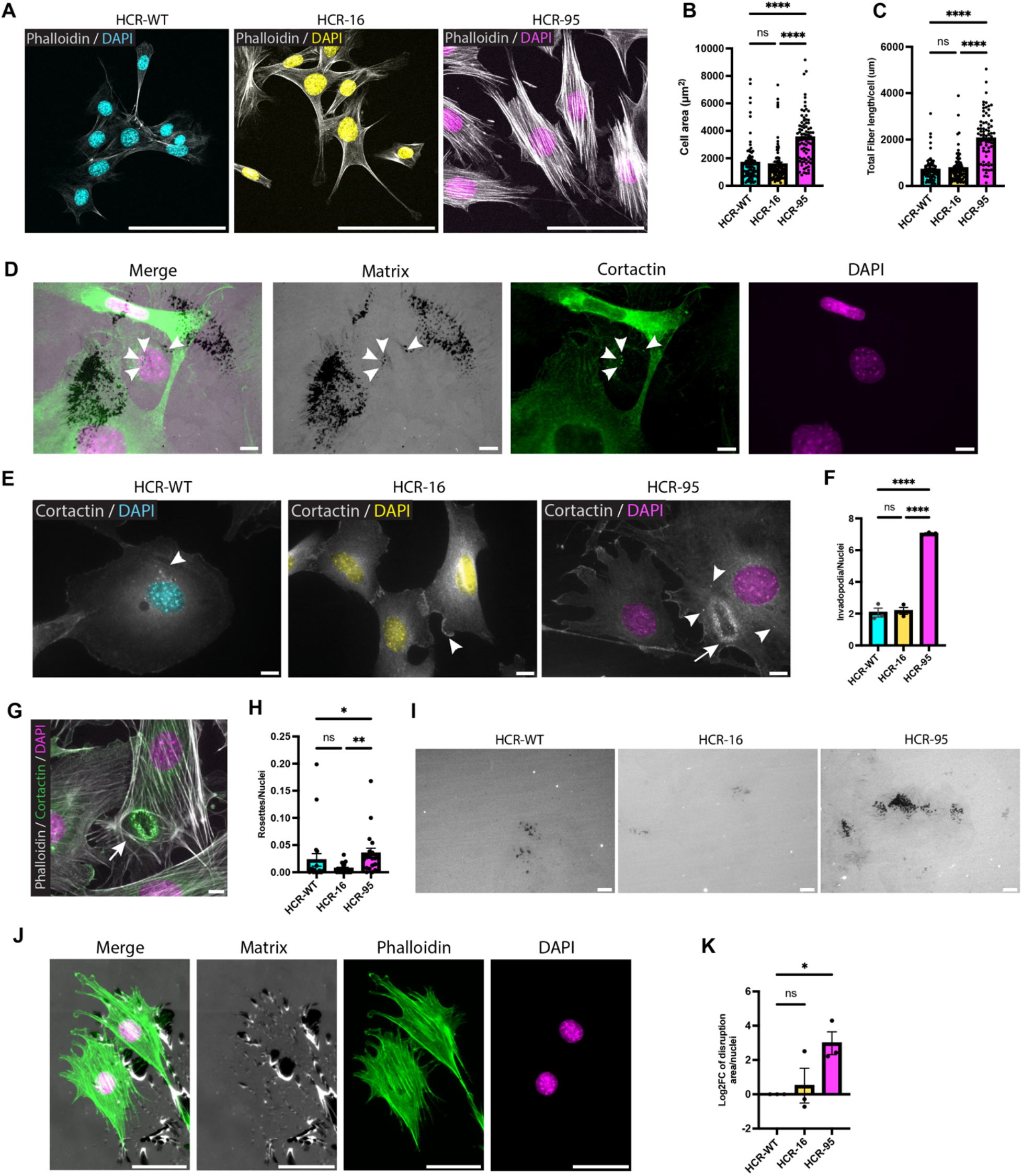
Comparing cellular morphology and invasive behaviors in HCR cell lines. **A,** Representative images of HCR-WT, HCR-16, and HCR-95 cells plated for 24 hours on gelatin-coated coverslips, stained with phalloidin (grey) and DAPI (pseudocolor: HCR-WT– cyan, HCR-16 – yellow, HCR-95 – magenta). **B,** Quantification of cell area measurements on phalloidin-stained images (n= 81-82cells/condition). **C,** Quantification of total F-actin fiber length per cell on phalloidin-stained images (n=75 cells/condition). **D,** Example of invadopodia structures overlying areas of fluorescent matrix degradation in HCR-95 cells (matrix – grey, degradation puncta - black, cortactin – green, DAPI – magenta pseudocolor). Arrowheads indicate active invadopodia. **E,** Representative images of each cell line stained with cortactin (grey), F-actin (not shown to enable better visualization of puncta only) and DAPI (pseudocolor: HCR-WT– cyan, HCR-16 – yellow, HCR-95 – magenta) to identify F-actin+/ cortactin+ puncta indicating individual invadopodia structures (arrowheads) and rosette invadopodia superstructure in HCR-95 (arrow with tail). **F,** Quantification of F-actin+/cortactin+ invadopodia structures in HCR cells (n=73 40x fields of view (FOV) across 3 biological replicates, points are average invadopodia/nuclei/FOV). **G,** Representative image of invadopodia rosettes (arrow with tail) seen in HCR-95 cells by staining with phalloidin (grey), Cortactin (green), and DAPI (magenta). H, Quantification of rosette number per nuclei, from live imaging of LifeAct transfected HCR cell lines (n=19-20 20x FOV across 2 biological replicates) **I,** Representative images of matrix degradation by each HCR cell line, seen as black puncta in regions where fluorescent gelatin plate coating was enzymatically degraded by invadopodia of overlying cells, **J,** Areas of mechanical matrix disruption below stress fiber adhesion points in HCR-95 (matrix – grey, degradation puncta - black, cortactin – green, DAPI – magenta pseudocolor). **K,** Quantification of matrix disruption area per nuclei (including both enzymatic degradation and mechanical disruption), displayed as log_2_ fold change. Statistical significance was determined by one-way ANOVA with Kruskal-Wallis multiple comparisons test (B, C, H),Tukey’s multiple comparisons test (F), or Holm-Sidak multiple comparisons test (K). ns, not significant; *p<0.05; **p<0.01; ***p<0.001; ****p<0.0001. Error bars are represented as mean±s.e.m. Scale bars: 100 *μm* (A), 50 *μm* (I), 10 *μm* (D, E, G, J).

To further examine real-time actin cytoskeletal dynamics, we transfected the cell lines with fluorescently labeled LifeAct plasmid and performed live cell imaging on monocultures (Supplemental movies 1-3). The dynamic formation of F-actin+ invadopodia puncta, as well as rosette invadopodia superstructures, were again observed more frequently in HCR-95 cells (Figure 3H). To test functional matrix degradation potential of the cell lines, we plated cells on fluorescent gelatin and measured area of matrix disruption. In addition to observing focal points of enzymatic degradation by invadopodia (Figure 3D, I), particularly in HCR-95 cells we observed areas of matrix disruption which appeared as if the gelatin had been forcefully ripped off the coverslip (Figure 3J). Interestingly, these areas often corresponded to focal adhesion complexes at the ends of stress fibers suggesting that contractions of these actin stress fibers may indeed be dislodging the matrix forcefully to generate mechanical matrix disruption in addition to enzymatic degradation. Overall, HCR-95 cells were able to disrupt significantly more matrix than HCR-WT or HCR-16 cells (Figure 3K). Taken together, these studies suggest HCR-95 cells are uniquely programmed for matrix breakdown and invasion.

### HER2 isoform induced bias in signaling programs

Next, we performed single cell RNA sequencing (scRNAseq) of each HER2BOW cell line (10,000 cells per genotype) to screen for transcriptional differences in each cell line. UMAP clustering and visualization revealed fairly uniform distribution of each genotype across clusters (Figure 4A) that included expression of luminal epithelial marker genes (*Epcam*, *Krt8*) and myoepithelial/basal marker genes (*Krt14* and *Vim*) (Figure 4B). This indicated that the majority of the cells in each genotype were in an epithelial-mesenchymal like state (45). We next identified genotype specific marker genes. We found genotype dependent changes in gene expression suggesting that unique cell signaling and cell behavior states exist in each genotype (Figure 4C). In particular, HCR-95 cells expressed several additional genes involved in EMT (*Lcn2*) and actin remodeling (*Acta2, Tgfb1i1*) (46,47). Furthermore, we found that key genes involved in myogenesis were upregulated in HCR-95 cells at the transcriptional level, including a modest increase in myocardin related transcription factor (*Mrtfa*), a critical mediator of myofibroblast development, which acts with its cofactor serum response factor (SRF) to turn on target genes involved in cytoskeletal remodeling, contractility, and motility, including *Tgfb1i1* (Figure 4D) (48). Interestingly, the rosettes described in Figure 3 have been shown to be driven by Hic5 (*Tgfb1i1)* and have roles in enhanced matrix disruption and invasion (46,49). We then validated these findings on the protein level and found that epithelial proteins EPCAM and E-Cadherin (CDH1) were downregulated in HCR-95, mesenchymal marker Vimentin (VIM) was upregulated, and multiple MRTFA targets including α-smooth muscle actin (ACTA2) and HIC5 (TGFB1I1) were upregulated, suggesting HCR-95 cells may indeed be undergoing an epithelial-myofibroblast transition (Fig 4E). Importantly, HCR-95 cells retained epithelial features while simultaneously expressing cytoskeletal and contractility-associated programs, consistent with a hybrid invasive cell state. Although whole cell expression of MRTFA was similar between isoforms, subcellular fractionation of cell lysates revealed increased nuclear:cytoplasmic ratio of MRTFA in HCR-95 cells, indicative of increased MRTFA transcriptional activity (47) (Figure 4F,G). We also confirmed these observations by visualizing MRTFA expression through immunostaining of the HCR cell lines, which showed increased nuclear translocation of MRTFA in HCR-95 cells (Figure 5A).

**Figure 4.**
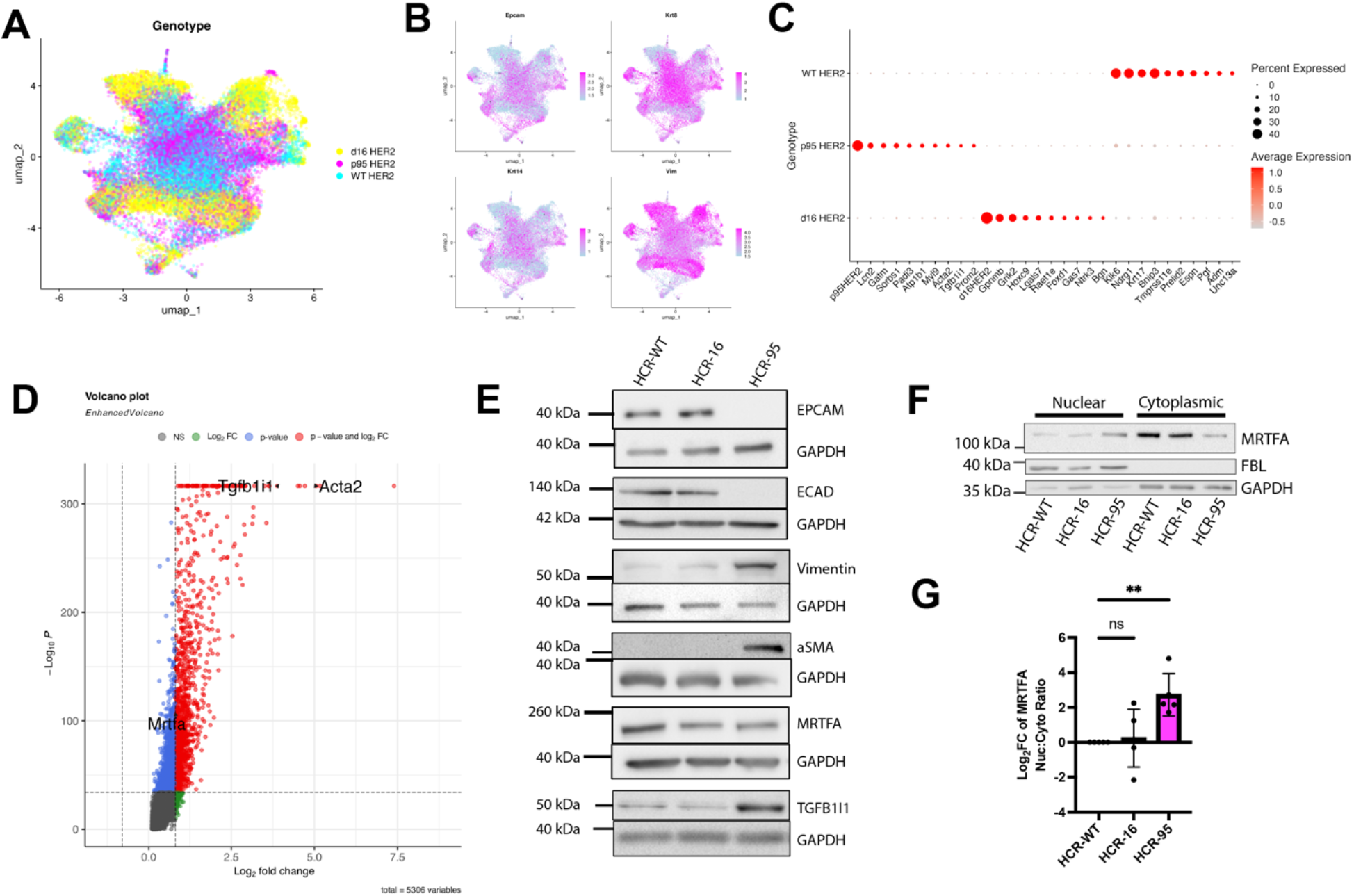
Single cell analysis of HER2 Crainbow cell lines. **A,** Single cell sequencing of HCR cell lines integrated and clustered by UMAP projection. **B,** Expression of epithelial and mesenchymal genes within UMAP clusters. **C,** Dot plot of genotype-specific markers in each cell line. **D,** Volcano plot of myogenic genes enriched in HCR-95 cells. **E,** Protein level expression of MRTFA and its gene targets in HCR cell lysates by western blot. **F,** Fractionation western blot distinguishing expression of MRTFA and fibrillarin in the nucleus and cytoplasm or HCR cell lines and **G,** quantification. Statistical significance was determined by one-way ANOVA with Holm-Sidak’s multiple comparisons test. ns, not significant; *p<0.05; **p<0.01; ***p<0.001; ****p<0.0001. Error bars are represented as mean±s.e.m.

**Figure 5.**
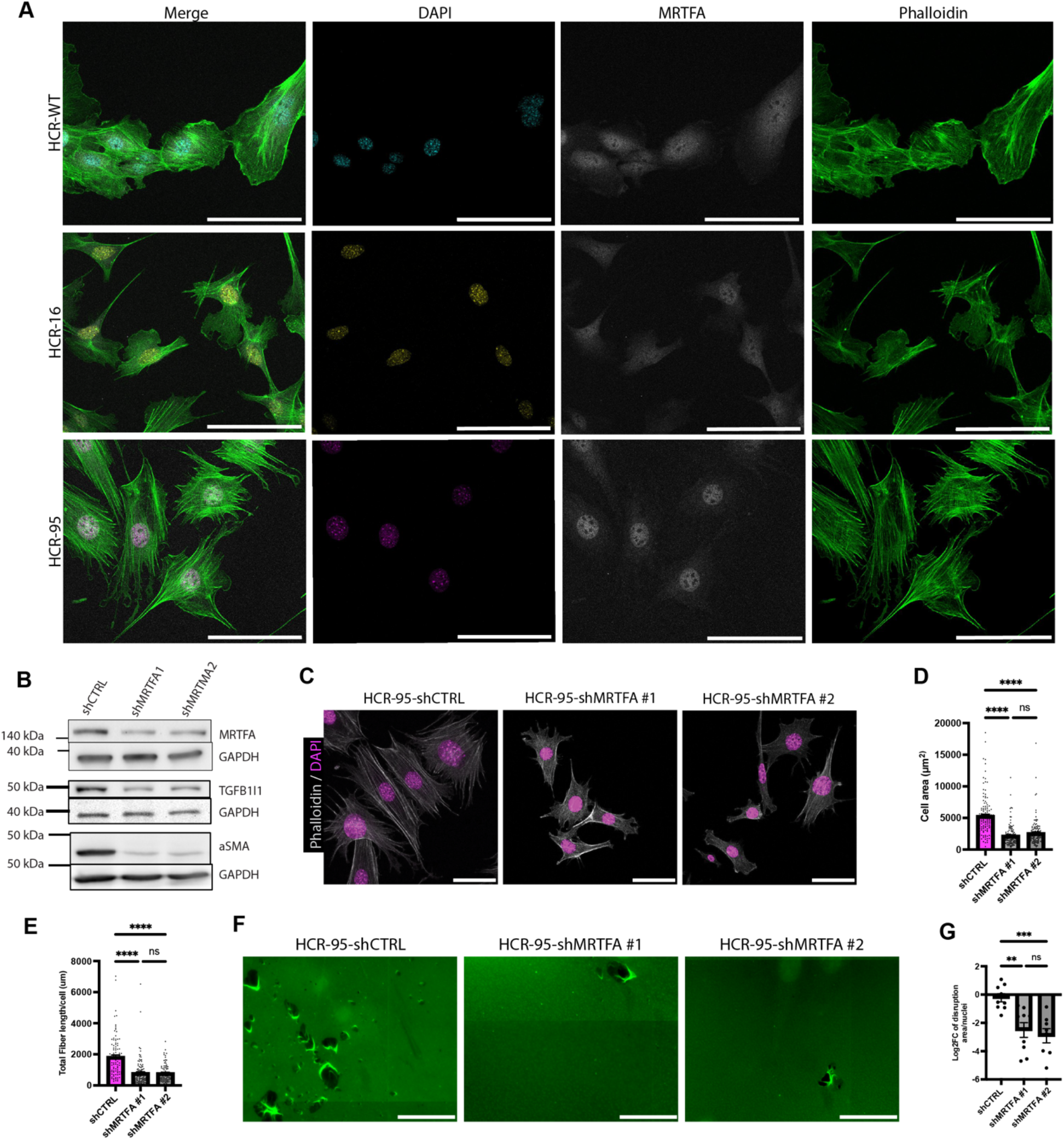
Short hairpin RNA knock-down of MRTFA leads to loss of invasive characteristics in HCR-95 cells. **A,** Representative images of HCR cell lines stained with phalloidin (green), anti-MRTFA (grey), and DAPI (pseudocolor: HCR-WT– cyan, HCR-16 – yellow, HCR-95 – magenta). **B,** Western blots of protein level expression of MRTFA and downstream targets aSMA and TGFB1I1 in HCR-95-shCTRL and HCR-95-shMRTFA cell lines. **C,** Representative images of HCR-95-shCTRL and HCR-95-shMRTFA cell lines stained with phalloidin (grey) and DAPI (pseudocolor: magenta) showing morphology and cytoskeletal organization. **D,** Quantification of cell area in HCR-95-shCTRL and HCR-95-shMRTFA cell lines on phalloidin stained images (n=128 cells/condition). **E,** Quantification of total F-actin fiber length/cell in HCR-95-shCTRL and HCR-95-shMRTFA cell lines (n**=**95-106 cells/condition). **F,** Representative images of matrix disruption by each HCR-95-shCTRL and HCR-95-shMRTFA cells, seen as black regions where fluorescent gelatin plate coating was degraded or ripped. **G,** Quantification of matrix disruption area per nuclei, displayed as log_2_ fold change. Statistical significance was determined by one-way ANOVA with Kruskal-Wallis multiple comparisons test (D, E) or Holm-Sidak’s multiple comparisons test (G). ns, not significant; *p<0.05; **p<0.01; ***p<0.001; ****p<0.0001. Error bars are represented as mean±s.e.m. Scale bars: 100 *μm* (A), 50 *μm* (E), 10 *μm* (H).

To determine if MRTFA expression was necessary for HCR-95 invasive phenotype and behavior, we utilized MRTFA-targeting shRNA to generate MRTFA-low HCR-95 cell lines (HCR-95-shMRTFA), and in parallel created a control cell line with a non-targeting shRNA (HCR-95-shCTRL). Proteins of MRTFA target genes, including ACTA2 and TGFB1I1, showed lower expression in the knockdowns (Figure 5B). Morphologically, MRTFA knockdown cells showed a loss of the spindle-like morphology seen in HCR-95-shCTRL and significant reduction in cell area (Figure 5C, D). F-actin stress fibers were also notably reduced in HCR-95-shMRTFA cells, showing shorter fibers that were less aligned in a parallel fashion, unlike in HCR-95-shCTRL (Figure 5E). Furthermore, the ability of the HCR-95-shMRTFA cells to disrupt extracellular matrix was significantly decreased (Figure 5F, G). TGFB1I1 was also among the top differentially expressed genes in HCR-95 cells and is a direct product of the MRTFA target gene transcriptional program (48). Therefore, we reasoned that TGFB1I1 could be an important downstream mediator of MRTFA in the p95-driven invasive phenotype. We confirmed that TGFB1I1 was indeed expressed at a much higher level in HCR-95 cells (Figure 4E, 6A), and that the protein localizes to focal adhesions at the ends of stress fibers (Figure 6B) as well as within invadopodia rosettes (Figure 6C), as expected from its described functions as a focal adhesion complex protein crucial for rosette formation (46,50,51).

**Figure 6.**
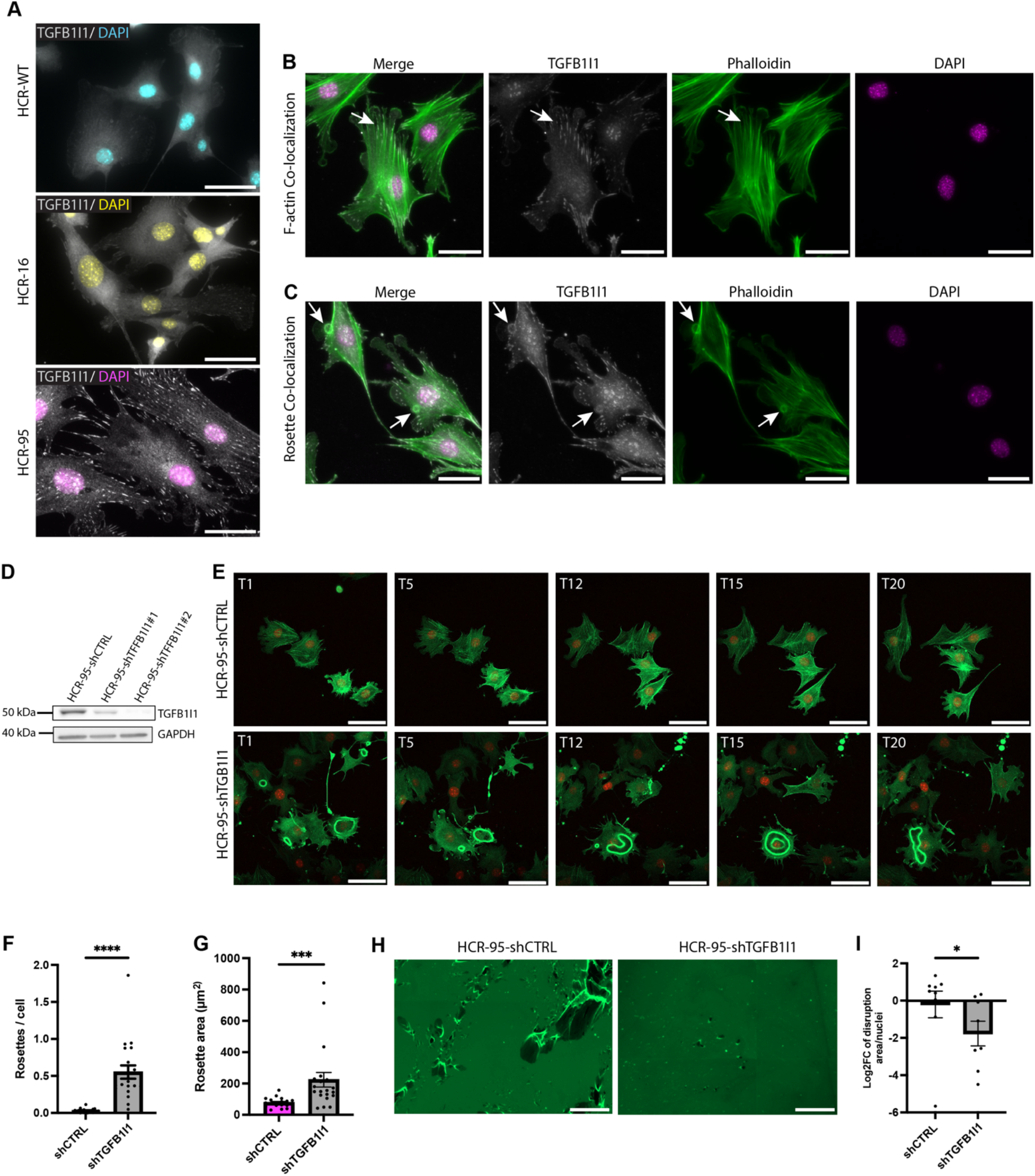
Knock-down of TGFB1I1 leads to loss of invasive characteristics in HCR-95 cells. **A,** Representative images of HCR cell lines stained with TGFB1I1 (grey) and DAPI (pseudocolor: HCR-WT– cyan, HCR-16 – yellow, HCR-95 – magenta). **B,** Representative images of HCR-95 stained with Phalloidin (green), anti-TGFB1I1 (grey), and DAPI (pseudocolor: magenta) to show TGFB1I1 co-expression with F-actin fibers (arrows). **C,** Representative images of HCR-95 stained with Phalloidin (green), TGFB1I1 (grey), and DAPI (pseudocolor: magenta) to show TGFB1I1 co-expression with rosette (arrows). **D,** Western blot of TGFB1I1 expression in two HCR-95-shTGFB1I1 knock down cell lines (shTGFB1I1 #1 and shTGFB1I1#2) and the vector control cell line (HCR-95-shCTRL). **E,** Timelapse live imaging of rosette formation using Lifeact in HCR-95-shCTRL and HCR-95-shTGFB1I1#2 showing formation of larger, more abnormally shaped and persistent F actin rings/rosettes upon TGFB1I1 knockdown (Supplemental Videos 4 and 6) . **F,** Quantification of rosette number in HCR-95-shCTRL and HCR-95-shTGFB1I1#2 from LifeAct imaging (n=18-20 20xFOV from 2 biological replicates; average rosette number/nuclei). **G,** Quantification of rosette area in HCR-95-shCTRL and HCR-95-shTGFB1I1#2 from LifeAct imaging (n=20 FOV from 2 biological replicates; average rosette area/nuclei). **H,** Representative images of matrix disruption by HCR-95-shCTRL and HCR-95-shTGFB1I1#2 cells, seen as black puncta regions where fluorescent gelatin plate coating was degraded or ripped. **I,** Quantification of matrix disruption area per nuclei, displayed as log_2_ fold change. Statistical significance was determined by Mann-Whitney test (F, G, I). ns, not significant; *p<0.05; **p<0.01; ***p<0.001; ****p<0.0001. Error bars are represented as mean±s.e.m. Scale bars: 50 *μm* (A, B, C, E, H).

Next, we wanted to determine the necessity of TGFB1I1 expression in HCR-95 rosette formation and functional degradation capability though shRNA knock-down (HCR-95-shTGFB1I1) (Figure 6D). Previous literature has noted that knocking down expression of TGFB1I1 in normal human mammary epithelial cells causes decreased invadopodia rosette formation and subsequently lower matrix degradation and invasion (46). Surprisingly, we noted an increase in invadopodia rosette formation in live imaging with Lifeact-transfected cells (Figure 6 E, F, Supplementary movies 4-8). However, these rosettes were abnormal in appearance, with much larger area and prolonged presence, rather than the transient appearance as seen in migrating HCR-95-shCTRL cells (Figure 6 E, G, Supplementary Movies 4-8). Furthermore, matrix disruption was decreased upon TGFB1I1 knockdown, suggesting that although cells are attempting to form invadopodia rosettes and they appear in increased number, with low TGFB1I1 these invadopodia structures are less functional and cannot drive successful matrix degradation and invasion (Figure 6H, I). Taken together, these results indicate that the MRTFA/TGFB1I1 signaling axis is essential for the invasive phenotype.

### HER2 isoform biased signaling at Y1139 drives “go” behaviors

We hypothesized that the p95-HER2 receptor selectively engages the MRTFA pathway through biased dependence on unique tyrosine autophosphorylation sites. Previous literature suggested Y1139 and Y1221/2 are sufficient to drive metastasis (“Go”) and proliferative (“Grow”) behaviors (40). Therefore, we performed co-transfections of control (EYFP) and EYFP-2a-HER2 isoforms into HEK cells along with an mTagBFP2-MRTFA fusion reporter for imaging nuclear MRTFA translocation (52). Furthermore, p95 mutants were made to test the necessity of Y1139 and Y1221/2 (p95^Y1139F^ and p95^Y1221/2F^).

Qualitatively, d16 and p95-HER2 promoted MRTFA translocation but p95 expression promoted the highest MRTFA translocation as compared to control (EYFP only). Furthermore, this effect was inhibited in the p95^Y1139F^ condition but not in the p95^Y1221/2F^ condition (Figure 7A). Quantitative analysis confirmed these observations (Figure 7B). To determine the necessity of Y1139 in the parental HCR-95 cell line, CRIPSR/Cas9 was used to make a Y1139F mutant stable cell line, herein called HCR-1139.

**Figure 7.**
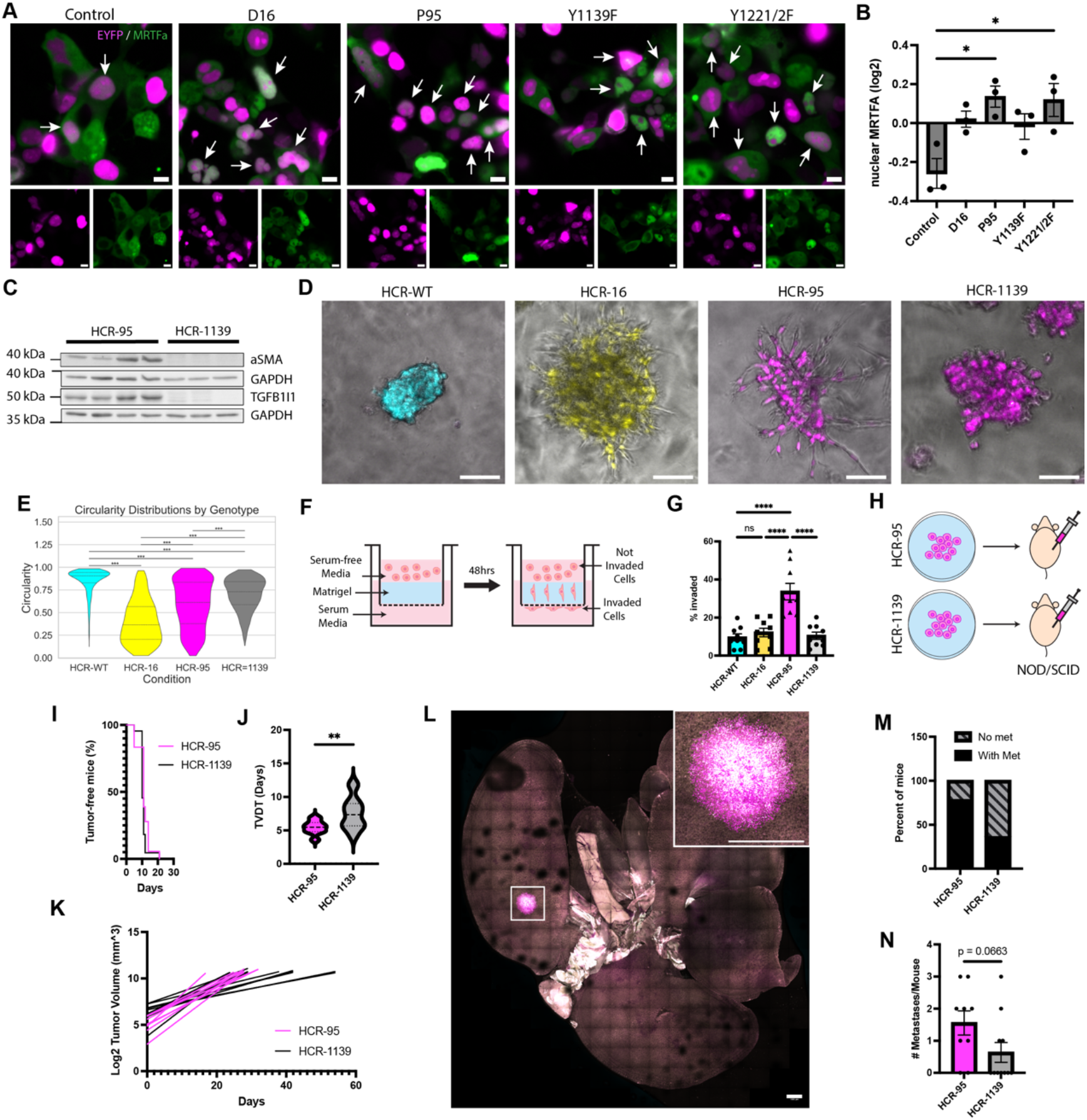
Functional selectivity of p95 receptor signaling through Y1139 promotes MRTFA activation and invasion. **A,** HEK cells co-transfected with either a control EYFP, EYFP-2a-d16 HER2, EYFP-2a-p95 HER2, EYFP-2a-p95^Y1139F^ HER2, or EYFP-2a-p95^Y1221/2^ HER2 (magenta) and an mTagBFP2-MRTFA fusion (green). **B,** Log2 fold change of MRTFA nuclear expression (n=3 biological replicates for a total range of 222 – 291 cells analyzed). **C,** Western blots of HCR-95 and HCR-1139 protein level expression of MRTFA targets ACTA2 and TGFB1I1 **D,** Representative images of 3D spheroid formation in Matrigel matrix at 96 hours. **E,** Quantification of sphere circularity distributions of each cell line. Experiment was performed 3 independent times, and all organoids were aggregated and compared (n=3 biological replicates for a total of 523-1144 spheres analyzed). **F,** Schematic of transwell invasion assay. **G,** Quantification by z-position of invaded cells as percentage of total population. **H,** Schematic of orthotopic transplants of HCR-95 and HCR-1139 into NOD/SCID mice. **I,** Tumor-free survival of orthotopic transplants of NOD/SCID mice with HCR-95 or the p95-Y1139F mutant (HCR-1139) cells. **J,** Linear regression of log2 transformed tumor volume measurements of HCR-95 (n=9) and HCR-1139 (n=11) transplants. **K,** Tumor volume doubling time of HCR-95 and HCR-1139 transplants. **L,** Clarified lungs of HCR-95 transplanted mouse with representative metastasis (inset zoom). **M,** Quantification of metastasis in HCR-95 and HCR-1139 transplanted mice by percentage of population with at least one metastatic lesion and **N,** number of metastases per mouse per condition. Statistical significance was determined by one-way ANOVA with Tukey’s multiple comparisons test (B, G) or Mann-Whitney test (J, N), or Anderson-Darling test with Dunn’s multiple comparisons test with Bonferroni correction (E). ns, not significant; *p<0.05; **p<0.01; ***p<0.001; ****p<0.0001. Error bars are represented as mean±s.e.m. Scale bars: 10 *μm* (A), 100 *μm* (D), 1000 *μm* (L).

Downregulation of MRTFA targets at the protein level (ACTA2, TGFB1I1) was confirmed in HCR-1139 relative to HCR-95 (Figure 7C). Using 3D Matrigel culture, we observed distinct morphological features of each cell line (Figure 7D). HCR-WT cells assembled as spherical clusters whereas HCR-16 and HCR-95 cells made stellate structures with cells branching away from a central sphere, indicative of highly invasive cells (53). Meanwhile, HCR-1139 cells created hybrid structures that were spherical with very little invasion away from the central sphere. Quantification of circularity distribution showed all conditions were significantly different from one another, with HCR-16 and HCR-95 scoring as the least circular and HCR-WT as the most circular (Figure 7E). In order to better quantify invasion potential of these cell lines, we leveraged a Boyden chamber assay in which we coated the microporous membrane in Matrigel matrix (Figure 7F) (54). We found that HCR-95 cells were significantly more invasive than HCR-WT and HCR-16 cells and that the Y1139F mutation inhibited invasive behavior (Figure 7G).

Finally, *in vivo* ‘go’ and ‘grow’ dynamics were assessed using orthotopic transplant into NOD/SCID mice with HCR-95 and HCR-1139 (Figure 7H). Tumor penetrance remained at 100% for both cell lines, however HCR-1139 transplants showed slower growth and had significantly higher doubling time (Figure 7I, J, K). To assess metastatic potential, the lungs of these animals were removed and clarified using FUnGI clarification, then imaged using confocal microscopy (Figure 7L) (55).

Metastases were quantified for both groups of animals, in which we found that a higher percentage of HCR-95 transplants had at least one metastasis present in their lungs (78%) as opposed to HCR-1139 transplants (35%) (Figure 7M). Furthermore, the number of metastases in HCR-95 transplanted animals trended higher, although not significant (Figure 7N). These results show that loss of function of Y1139 leads to phenotypically and functionally less invasive cells that are less likely to form metastases.

## Discussion

‘Go’ (invasion and metastasis) and ‘grow’ behaviors (proliferative signaling) are fundamental hallmarks of cancer shaping cancer progression (11). Yet, a major problem is that these behaviors need not occur linearly. For instance if invasion occurs prior to or in parallel to major proliferative advances, then cancer cells can disseminate and metastasize prior to detection (56,57). Previously we demonstrated that d16 HER2 tumors appear to progress linearly to metastasis, whereas p95 HER2 tumors progress to metastasis parallel to their initiation and growth at the primary tumor site (9). The goal of the current study was to determine how HER2 isoforms could promote these distinct pathways. Overall, we used several empirical observations to construct and build a summary model by which each isoform differentially promotes tumor cell behaviors (Figure 8). Although each genotype is capable of promoting any one of a number of hallmarks, like go or grow, the isoforms are nonetheless quantitively different.

**Figure 8.**
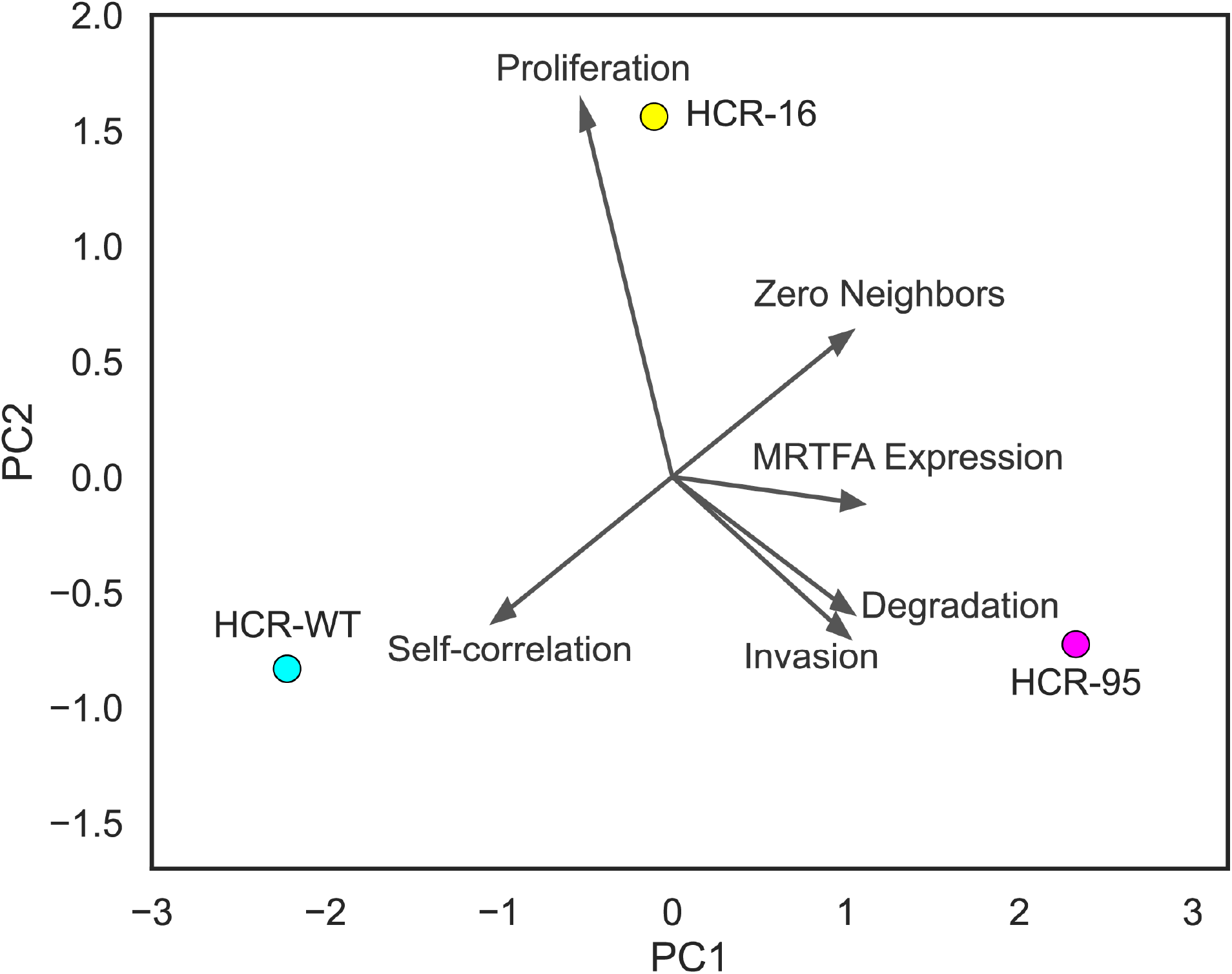
Summary of functionally selective behaviors engaged by HER2 isoforms. Principal component model representing key cell line phenotypes. Average values from six different assays were collected and scaled for PCA analysis: Proliferation = inverse tumor volume doubling time from *in vivo* transplantation, Self-correlation and Zero Neighbors = self-correlation and nearest neighbors kinematic motility analyses, MRTFA Expression = whole cell lysate western blot analysis, Invasion = Boyden chamber transwell invasion quantification, Degradation = fluorescent degradation quantification.

For instance, WT cells exhibit increased collective behavior (“Self-correlation”) and are much less likely to invade, degrade matrix, and activate MRTFA. In contrast, even though both d16 and p95 are each able to invade/degrade, d16 is biased toward increased proliferation compared to p95, which is most biased toward MRTFA activation and invasion.

A major question was whether the phenotypes we observed could be due to receptor-intrinsic isoform signaling differences, or if they were the result of adaptions *in vivo* that led to cell lines with distinct phenotypes. To answer this question, we performed single cell analyses and found that cell lines showed many similarities in epithelial and mesenchymal gene expression but HCR-95 cells were enriched in genes associated with a myogenic-like transition – for instance *Acta2*, *Tgb1i1*, and *Mrtfa* as discussed below. Importantly, we confirmed this data by showing that transient transfection of the p95-HER2 receptor into HEK cells was able to promote biased MRTFA nuclear translocation, compared to more modest effects of d16-HER2 on HEK nuclear MRTFA localization. This suggests that receptor-intrinsic behaviors of each isoform biases MRTFA signaling, yet we cannot entirely exclude other potential cell line-specific effects that only a deeper analysis of other HER2 Crainbow cell clones could provide.

We also provide a unique source of HCR cell lines that retained important features demonstrated in the HER2-BOW model. This includes isoform-specific tumor cell properties that promote invasion and dissemination (p95-HER2/HCR-95) and proliferation (d16-HER2/HCR-16). Each cell line is capable of being cultured for at least 30 passages while also retaining their expression of human isoforms and appropriate fluorescent barcodes for live cell analyses. *In vitro* characterization of the cell lines points to HCR-95 being functionally more invasive than both HCR-WT and HCR-16. Moreover, HCR cell phenotypes *in vitro* can also be replicated across multiple conditions including *in vivo* orthoptic transplant. Together, this confirms previous work in the autochthonous HER2 Crainbow model which demonstrated how differences in early dissemination and proliferative potential could be encoded by each isoform. Furthermore, these findings confirm the presence of heritable phenotypes that promote heterogeneity in cell behaviors “capable of breeding true” (58).

The discovery of a highly active MRTFA-driven transcriptional program was unexpected and potentially provides a unifying mechanism explaining how p95-HER2 exerts its effect on tumor progression. MRTFA is a transcription factor that promotes cell motility, migration, invasion, and cell state plasticity (59). It is downstream of Rho-driven actin cytoskeletal rearrangements that promote its release from G-actin and enable nuclear translocation and both SRF-dependent and independent signaling, and its expression increases metastatic potential and is correlated with decreased breast cancer survival (60–63). One important downstream MRTFA target is TGFB1I1, which promotes a contractile phenotype, inhibits cell proliferation, and is co-expressed with alpha smooth muscle actin (SMA) (64). Furthermore, MRTFA also promotes an epithelial to myofibroblast transition (65). Interestingly, our scRNAseq and protein validation each indicate that both ACTA2 and TGFB1I1 are upregulated in HCR-95 cells. This suggests that HCR-95 cells are not only contractile but could also have undergone an epithelial to myofibroblast like transition, or as previously reported this could be similar to a luminal to basal-like transition (66,67). Furthermore, recent reports in breast cancer cell models show MRTFA driven increases in cell migration, invasion, and metastatic potential (59,68). Similarly, we show that each of these behaviors occurs with increased frequency in HCR-95 cells and we demonstrate that knockdown of MRTFA or TGFB1I1 can inhibit these behaviors. Lastly, our data provide an important link to recent reports illustrating that p95-HER2 promotes immune evasive microenvironments, as MRTFA can also promote PDL1 expression (26,69,70). Thus, our data support a model by which HER2 can directly promote metastatic potential by engaging the MRTFA-TGFB1I1 pathway.

Our study is the first one to report how HER2 isoforms regulate invasive protrusions of tumors cells. Invadopodia are actin-rich protrusion that mediate matrix degradation and facilitate dissemination by degrading ECM during invasion and intravasation into the blood vessels (43,44). Our work points towards a novel signaling pathway controlling invadopodium formation in HER2 tumors driven by p95, the MRTFA/TGF1I1 axes. Interestingly the protrusive plasticity observed in p95 cells with increased invadopodia protrusions and invadopodia rosettes strongly support the highly motile and invasive nature of these cells. Moreover, MRTFA activity is regulated through actin polymerization-dependent nuclear translocation, further suggesting a potential link between p95-HER2 signaling and cytoskeletal remodeling pathways. Our results point towards the MRTFA/TGFB1I1 pathway as a potential target to prevent further dissemination from HER2 tumor cells in patients that express p95-HER2.

Our data also demonstrate how functional selectivity could explain differences between HER2 isoforms. Functional selectivity can occur via ligands that promote distinct conformational changes in the receptor or even through changes in the expression of downstream effectors within a particular cell (35,71). Many times, this is also associated with unique phosphorylation of the C-terminal phosphobarcode in a receptor, that enhances the affinity for downstream effectors (72,73). The result is that the same receptor can exist in multiple states, each capable of being biased toward a particular signaling pathway given the right ligand or context. Similarly, functional selective signaling has been shown to occur in RTKs (36,37). HER2 alone has well-studied phosphotyrosine sites that are autophosphorylated in the Carboxyl-tail, referred to as YA (human Y1023), YB (human Y1139), YC (human Y1201), YD (human Y1221/2), and YE (human Y1248). Previous work in mouse models has shown that reconstitution of YB or YD in a tyrosine-dead rat Neu transgenic are both sufficient to initiate and promote tumorigenesis (74). Yet, YB had increased tumor latency and increased metastatic potential.

Our work confirms these findings and extends them by showing how loss of signaling of YB could prevent MRTFA mediated cell motility, invasion, and metastatic behavior.

There are limitations to our current study. A central limitation is that all in vivo experiments were conducted in NOD/SCID hosts, which was necessary to engraft humanized HER2 isoform–expressing cells and to isolate receptor-intrinsic contributions to growth and dissemination, but it constrains interpretation. The metastatic phenotypes reflect tumor-cell-intrinsic invasive capacity in the absence of adaptive immune surveillance, and the absolute metastatic burden in an immune-competent host may differ. Our discussion of p95-associated immune-evasive programs, including JAK/STAT and IL-6 signaling and MRTFA-driven PD-L1 induction, is drawn from prior work (25,75–77) and is intended to motivate future study rather than as a conclusion supported by the present data; as our current model cannot test whether the MRTFA/TGFB1I1 axis shapes the immune microenvironment. Addressing this will require competitive transplantation of fluorescently barcoded isoform-pure lines into immune-competent or humanized hosts which would preserve the direct isoform-versus-isoform comparison that is the strength of this model while restoring the immune context in which p95-driven progression and therapeutic resistance ultimately occur. Additionally, we focus on the characterization of three HER2 Crainbow cell lines. We do not assess the possibility that each cell line could represent a distinct clone whose behavior may be distinct. Although this is a known issue with any cell line model, future work could assess WT, d16, and p95 cell lines from tumors of different age and size. Also left unresolved is how HER2 can couple to MRTFA and whether or not the remaining tyrosine residues in the C-tail of HER2 play additional roles in this signaling pathway. Lastly, determining whether p95-driven MRTFA signaling is sensitive to RTKi, p95-specific monoclonal antibody therapy, or MRTFA inhibition could provide insight into inhibiting metastatic potential (26,78,79).

A central limitation is that all in vivo experiments were conducted in NOD/SCID hosts, which was necessary to engraft humanized HER2 isoform–expressing cells and to isolate receptor-intrinsic contributions to growth and dissemination, but it constrains interpretation. First, the metastatic phenotypes reflect tumor-cell-intrinsic invasive capacity in the absence of adaptive immune surveillance, and the absolute metastatic burden in an immune-competent host may differ. Our discussion of p95-associated immune-evasive programs, including JAK/STAT and IL-6 signaling and MRTFA-driven PD-L1 induction, is drawn from prior work (25,26,69,70) and is intended to motivate future study rather than as a conclusion supported by the present data; as our current model cannot test whether the MRTFA/TGFB1I1 axis shapes the immune microenvironment. Addressing this will require competitive transplantation of fluorescently barcoded isoform-pure lines into immune-competent or humanized hosts which would preserve the direct isoform-versus-isoform comparison that is the strength of this model while restoring the immune context in which p95-driven progression and therapeutic resistance ultimately occur.

Strikingly, we found that HCR-95 biases signaling toward an invasive phenotype through Y1139-dependent activation of an MRTFA-dependent signaling pathway (Figure 8). Together this suggests that the HER2 Crainbow cell lines retain biased “grow” behavior (WT and d16) and “go” behavior (p95) that we previously reported *in vivo* (9,10). These data provide a working model whereby functionally selective HER2 isoform signaling can be engaged to preferentially drive ‘go’ behavior prior to or in parallel with ‘grow’ behavior, thereby implicating naturally occurring isoforms of HER2 as a potential source of molecular adaptation during progression to treatment-resistant breast cancer.

## Materials and Methods

### Establishing HCR cell lines

#### Establishing & Maintaining Luminal Crainbow HER2 (HCR) cell lines

Protocol for establishing HCR cell lines was adapted from Ginzel *et al*. (9) Overall mammary tumor burden of HER2BOW mice aged between 15-20 weeks were characterized using the Zeiss Axio Zoom fluorescent stereoscope after sacrifice. Tumors of interest were resected and mechanically digested with scalpels in 1mL PBS in a 10cm dish. Tumor digests were pipetted into 15mL tubes with 5mL of RPMI media (Gibco 11875093) with 1% antibiotic/antimycotic (Gibco 15240096) and 2mg/mL Collagenase A (Millipore Sigma 10103578001), then incubated at 37C for up to 2 hours, with gentle inversion every 15 minutes. Digest samples were spun down at 150 x g for 5min and resuspended in 5mL of 1:100 DNase I (Thermo Fisher Scientific 18047019):RPMI solution. This was followed by RBC lysis (Millipore Sigma R7757), PBS quench, and centrifugation. Pellets were resuspended and counted using the Bio-Rad TC20 automated cell counter. Cells were seeded and maintained in DMEM (Gibco 11965092) with 10% fetal bovine serum (Gibco 16000044) and 1% antibiotic/antimycotic (DMEM 10%) until single genotype cultures were achieved.

#### Creation of shRNA-knockdown cell lines

Lentiviral transfer plasmids (listed below) and plasmids encoding VSV-G, Gag-Pol, Rev, and Tat genes were transiently transfected into HEK293T cells (ATCC, CRL-11268) with Lipofectamine™ 2000 (Thermo Fisher Scientific 11668027). After incubation at 37C for 72h for lentiviral packaging, virus-containing supernatant was collected. Cell lines were infected with virus in DMEM 10% with 5ug/ml of polybrene for 48h, after which virus containing media was removed and cells washed with PBS. Lentiviral shRNA infected cell lines were subjected to further culture in selection media (5 ug/ml puromycin) for at least 1 week after cells in control non-infected wells had all died. Lentiviral shRNA infected cell lines were analyzed by western blotting to confirm target knockdown. Full western blots shown in supplemental figures.

shRNA lentiviral plasmids:

shCTRL – TRC Lentiviral Non-targeting Control

shRNA (Horizon, RHS6848)

shMRTFA-1 (Albert Einstein College of Medicine

shRNA core facility Cat# RMM3981-97074500; sequence CCCACTCAGGTTCTTTCTCAA)

shMRTFA-2 (Albert Einstein College of Medicine

shRNA core facility Cat# RMM3981-97074502; sequence CAGGTAAATTACCCAAAGGTA)

shTGFB1I1-1 (Albert Einstein College of Medicine shRNA core facility Cat# RMM3981-97063816; sequence AGACCACTTCACCTGCACATT)

shTGFB1I1-2 (Albert Einstein College of Medicine shRNA core facility Cat# RMM3981-97063818; sequence GACCGTTTGCTTCAGGAACTT)

#### Creation of HCR-1139 phosphomutant cell line

We electroporated HCR-95 cells with a complex of Cas9 protein and sgRNA (Integrated DNA Technologies), and a single strand DNA oligo as the HR template using the NEON electroporator (Thermo Fisher Scientific). The sgRNA target sequence is gctggttcacatattcaggc, and the HR template sequence is GGGCCCTCTCGGGGCGAAGGGGGCTGGGGCCGAACATCTGGCTGGTTCACGAATTCCGGTTGGGGGCTGCAGGTCAGGGGGGCAACGTAGCCATCAGTCTCAGAGGGC. The cells were plated onto 96 well plates at a density of 0.7 cell per well 2 days after electroporation. When the clones grew back, they were screened by genomic PCR and EcoRI digestion. The PCR primers used are Fw-tctgaattctcccgcatggc and Rev-gggtgtcaagtactcggggt. Positive clones were confirmed by Sanger sequencing (Supplemental Figure S21).

### Protein-level expression and localization in vitro assays

#### IHC staining of cell lines and transplant tumors

Immunohistochemistry (IHC) was performed according to previously reported procedures (PMID: 26157038). Cell pellets were prepared in HistoGel (Thermo Fisher Scientific) and embedded in formalin-fixed paraffin-embedded (FFPE) blocks. Four-micron sections of FFPE cell pellets or tissues were cut and mounted onto Superfrost Plus microscope slides (Thermo Fisher Scientific). Sections were deparaffinized, subjected to antigen retrieval in CC1 buffer (pH 9.0, 95°C; Roche), blocked for endogenous peroxidase activity, and subsequently incubated with primary antibodies. Chromogenic detection was performed using the VECTASTAIN Elite ABC-HRP kit (Vector Laboratories), followed by counterstaining with hematoxylin. Whole-slide digital images were acquired on the Aperio Leica ScanScope XT slide scanner (Leica Biosystems, Richmond, IL) using a 40× objective lens. Primary antibodies used for this study are anti-HER2 C-terminus (clone CB11, Abcam) and anti-HER2 N-terminus (clone D8F12, Cell Signaling Technologies).

#### Immunofluorescent staining for cell morphology, actin cytoskeleton, invadopodia, and MRTFA localization

Cells were seeded at a density of 15,000 cells/well in 24 well plates on glass coverslips pre-treated with gelatin (Sigma Aldrich 9000-70-8) for 20 minutes and then 0.5% glutaraldehyde for 40 minutes, followed by 3x PBS washes prior to cell seeding. Cells were incubated for 24 hours in DMEM (Gibco ) with 10% FBS (Gemini Bio) and 1% penicillin/streptomycin (Gibco), then fixed with 4% paraformaldehyde in PBS for 10min, and washed 3x with PBS. They were incubated in permeabilization/blocking buffer (0.2% Triton X-100, 4% BSA in PBS) for 10min, washed 3X with PBS, then and incubated for 40 minutes at room temperature with the primary antibodies listed below, in 4% BSA/PBS. Coverslips were then rinsed 2× in PBS and incubated with secondary antibodies listed below and DAPI in 4% BSA/PBS for 30min at room temperature. They were then rinsed 2x in PBS and mounted on slides using Invitrogen Fluoromount-G polymerizing medium. Images were captured at the Microscopy and Advanced Bioimaging CoRE of the Icahn School of Medicine at Mount Sinai, with Zen Blue software on a Zeiss LSM980 microscope (Carl Zeiss Microscopy GmbH, Jena, Germany) equipped with GaAsP PMT detectors (Hamamatsu Photonics, Shizuoka, Japan) in confocal mode, at 40x magnification (Carl Zeiss Microscopy GmbH, Jena, Germany). For immunofluorescence on cells, primary antibodies were diluted in 4% BSA in PBS and added to coverslips for 40minutes at RT: Rabbit monoclonal [EP1922Y] to Cortactin (abcam, ab81208; 1:200), HIC5 Polyclonal Antibody (Invitrogen PIPA5117133; 1:500), MKL1/MRTF-A (E2V2I) Rabbit mAb (Cell Signaling #97109;l 1:200). Phalloidin and AlexaFluor labelled-secondary antibodies were diluted in 4% BSA in PBS from 1:200-1:400, and DAPI was used at 1:10,000 dilution and added to the secondary incubation: Donkey anti-Rabbit IgG (H+L) Alexa Fluor™ 546 (Invitrogen, A10040), Donkey anti-Rabbit IgG (H+L) Alexa Fluor™ 647 (Invitrogen, A31573), Donkey anti-Rabbit IgG (H+L) Alexa Fluor™ 488 (Invitrogen, A21206), Alexa Fluor™ 647 Phalloidin (Invitrogen, A22287), DAPI (4’,6-Diamidino-2-Phenylindole, Dihydrochloride) (Thermofisher, D1306).

#### Morphology and actin fiber analysis

Cell area measurements were analyzed in FIJI by using thresholding on a phalloidin-stained image to create a mask of the whole cell. Area of at least 75 cells in total per condition from a minimum of 15 different 40x FOVs across 2 independent experiments were quantified.

Analysis of F-actin filaments in phalloidin-stained specimens was performed using the Matlab algorithm developed and described by Rogge *et al*. which segments and analyzes cytoskeletal stress fibers (80).

Default standard parameters were utilized for the analysis.

Fiber analysis of at least 75 cells in total per condition from a minimum of 15 different 40x FOVs across at least 2 independent experiments was performed to compare total F-actin fiber length per cell.

#### Invadopodia Quantification

Immunofluorescence staining for cortactin and phalloidin, and image acquisition was performed as described above, and total number of cortactin and phalloidin co-positive puncta per cell was manually annotated and counted for at least 70 different 40x FOVs in total (minimum of 500 cells), across 3 independent experiments.

#### MRTFA fluorescent reporter assay

12-well Mattek glass bottom plates (Mattek P12G-0-14-F) were coated with 5ug/mL Fibronectin solution and incubated at room temperature for 15 minutes. Fibronectin solution was removed and replaced with DMEM 10%. HEK-293T cells were then seeded at 100,000 cells per well and allowed to incubate overnight. Cells were transfected with 1ug of EYFP1-tagged HER2 receptor (PB-CMV-EYFP1-2a-P95, PB-CMV-EYFP1-2a-P95-Y1139F, PB-CMV-EYFP1-2a-P95-Y1221/2F, PB-CMV-EYFP1-2a-D16) or control plasmid (PB-CMV-EYFP1) along with 1ug of an BFP-tagged MRTFA plasmid (pcDNA3.1(+)-mTagBFP2-MRTF-A (Addgene #182061)) using Lipofectamine 3000 (Thermo Fisher #L3000015 ). 24 hours post-transfection, media in plates were replaced with DMEM 10%. 48 hours post-transfection, cells were imaged live with the Zeiss 880 confocal microscope at 40x magnification (n=4 ROI/well). Dual-positive YFP/BFP nuclei were segmented in IMARIS imagining software. Nuclear BFP expression was log2 transformed, then averaged for each condition. Following this, expression for all conditions within each biological replicate. Finally, the log2 for each condition was divided by the log2 average for each biological replicate to determine how nuclear BFP expression of each condition compared to the biological replicate average nuclear BFP expression.

#### Western blots

Cells were lysed in RIPA buffer (25mM Tris HCl pH 7.5, 150Mm NaCl, 1% NP-40, 1% sodium deoxycholate and 0.1% SDS) with protease inhibitors (cOmplete™ Protease Inhibitor Cocktail, Roche ) and phosphatase inhibitors (phosSTOP™, Roche ), sonicated, and quantified using the Pierce™ BCA Protein Assay (Thermo Fisher). 10-60 micrograms (depending on the experiment) of protein was incubated at 95 °C for 5min in 4× Laemmli (Bio-Rad) with 10% beta mercaptoethanol and loaded onto a 10% SDS–PAGE gel. Proteins were transferred onto a nitrocellulose membrane (Bio-Rad) using the Bio Rad Turbo Transfer system . Ponceau S solution (Sigma) was used to validate even transfer across the membrane. Membranes were blocked with 5% BSA (Thermo Fisher) in TBS-T and probed with primary antibody overnight. Membranes were then washed 3x in TBS-T and incubated with the corresponding HRP-conjugated secondary antibody (1:5,000-1:15,000 dilution) for 2h at room temperature. Blots were again washed 3x with TBS-T, and signals were acquired and quantified with the ChemiDoc™ Touch Imaging System (Bio Rad). To probe for multiple proteins sequentially, blots were then incubated for 15 minutes with Restore™ PLUS Western Blot Stripping Buffer (Thermo Fisher), washed 3x in TBS-T and re-blocked with 5% BSA for 1h at RT before staining with another primary antibody. Full western blots shown in supplemental figures.

Primary antibodies used for westerns were as follows:

HIC5 Polyclonal Antibody (Invitrogen, PIPA5117133, 1:1500) MKL1/MRTF-A (E2V2I) Rabbit mAb (Cell Signaling #97109. 1:1000) GAPDH (D16H11) XP® Rabbit mAb (Cell Signaling #5174, 1:1000) alpha-Smooth Muscle Actin (D4K9N) Rabbit Monoclonal Antibody (Cell Signaling #19245, 1:1000) EpCAM (E6V8Y) Rabbit Monoclonal Antibody (Cell Signaling #93790, 1:1000) E-Cadherin (24E10) Rabbit Monoclonal Antibody (Cell Signaling #3195, 1:1000) Vimentin Antibody (R&D Systems, MAB2105, 1:1000) HRP-conjugated secondaries were diluted 1:5,000-1:10,000 in 5%BSA in 1XTBST and incubated with blots for 2h at RT after primary removal and 3x PBS washes for 10 minutes with shaking. For some blots, HRP-conjugated β-Actin was used for loading control, and this was incubated for 1h at RT at at dilution of 1:15,000 in 5%BSA in 1XTBST.

Anti-β-Actin−Peroxidase antibody (Sigma Aldrich, A3854; 1:15,000) Goat anti-Rabbit IgG (H+L) Secondary Antibody, HRP (ThermoFisher, 31460, 1:10,000) Goat anti-Mouse IgG (H+L) Secondary Antibody, HRP 31431 (ThermoFisher, 31460, 1:10,000) Anti-rat IgG, HRP-linked Antibody (Cell Signaling #7077, 1:5000)

Fractionation

Fractionation of fresh cell pellets into nuclear and cytoplasmic fractions was performed using NE-PER Nuclear and Cytoplasmic Extraction Reagents Kit (Thermo Scientific, #78833) according to manufacturer’s protocol.

### Integrated single cell RNA sequencing analysis

10,000 cells per line were captured using 10X Genomics universal 5’ single cell kit paired with the 10X Chromium instrument. Cell capture and library prep were performed per 10X Genomics published protocol and sequenced on Illumina NovaSeq 6000 (MedGenome Labs Ltd., Foster City, CA).

Sequencing reads were processed using CellRanger. Raw 10X Genomics gene-cell barcode matrices were converted to objects in Seurat (81). Cells were filtered based on total UMI, total features and mitochondrial, ribosomal, and hemoglobin gene counts. The percentage of reads mapping to transgene sequences (WT HER2, p95 HER2, d16 HER2) was calculated per cell and regressed out during data scaling to remove any effects of transgene expression. Principal component analysis (PCA) was performed on scaled data, and the top 30 principal components were retained. Integration across conditions was performed using Canonical Correlation Analysis (CCA) via IntegrateLayers (31178118). Nearest-neighbor graphs were constructed on the CCA-corrected embedding (dims 1–30) and cells were clustered using the Louvain algorithm at multiple resolutions (0.05–1.0); a final resolution of 0.4 was selected for downstream analysis. Differentially expressed genes between genotypes were identified using FindMarkers with default Wilcoxon rank-sum testing, retaining only positive markers expressed in at least 10% of cells. Results were visualized using dot plots, feature plots, and volcano plots generated with EnhancedVolcano (82). All analysis was performed in R using the Seurat framework. Sequencing data is available upon request.

### Cell function in vitro assays

#### Proliferation assay

HCR cells were seeded into a 96-well plate (GenClone 25-109MP) at 3000 cells/well in 10% DMEM. Outer wells were avoided due to edge effects. Starting 4-6 hours post-seed, wells were imaged at 5x once a day for 6 days using the Zeiss Cell Discoverer 7. Images were stitched in Zen Blue, segmented using IMARIS imaging software, then quantified using ‘R’ and GraphPad Prism.

#### Motility and kinematics assay

Wells of a Tissue culture-treated 96-well plate (GenClone 25-109MP) were coated with 5ug/mL of Fibronectin (R&D Systems 1030-FN-01M), Poly-D-Lysine (poly d lysine), or left uncoated (control -Plastic) for 15 minutes, then washed 3x with PBS. HCR cells were seeded in an equally mixed culture at serial dilutions from 10,000 cells/well to 313 cells/well in 10% DMEM. Starting at 4 hours post-seed, plates were imaged at 10x magnification every 10 minutes for 16 hours using the Zeiss Cell Discoverer 7. Cell motility was evaluated using three outcomes, namely, average speed, persistence, and mean squared displacement (MSD) slope. Cell-cell neighborhoods were evaluated using self-correlation and number of neighbors within 50 mm. See supplementary methods for additional information.

#### 3D Spheroid assay

40uL of Matrigel matrix (Corning 354234) per well was added to a round-bottom 96-well plate (Corning 3799) and incubated for 30 minutes at 37C to allow for polymerization. Next, HCR monocultures were seeded at 40,000 cells/well in 10% DMEM media. Immediately following cell addition, 10uL of addition Matrigel was added to each well. After 96 hours, spheres were imaged using the Zeiss 880 confocal microscope, segmented in IMARIS image analysis software, and quantified for circularity using Python script.

#### Transwell Invasion Assay

8um pore transwell inserts (Corning 3422) were coated with 100uL 1:8 diluted Growth Factor-Reduced Matrigel (Corning 356231) and incubated at 37C for 30 minutes. HCR monocultures were seeded at total 25,000 cells/well in either DMEM + 10% FBS (control) or DMEM serum-free (experimental) media.

Inserts were placed into wells with DMEM + 10% FBS media and incubated at 37C for 48 hours. At assay endpoint, inserts were imaged using z-position imaging at 1 mm steps using the Zeiss 880 confocal microscope. Cells were segmented by z-position using IMARIS image analysis software, then quantified in GraphPad Prism.

#### Matrix Disruption Assay

Glass coverslips were coated with 1mg/ml AlexaFluor-488 or AlexaFluor-546 labelled gelatin matrix (Thermo Fisher) as previously described (83). Cells were plated on coverslips at a density of 15,000 cells/well in 24 well plates, and incubated for 24 hours at 37C in DMEM (Gibco) with 10% FBS (Gemini Bio) and 1% penicillin/streptomycin (Gibco). Cells were then fixed with 4% PFA in PBS for 10min and washed 3X with PBS, then stained with AlexFluor-647 conjugated Phalloidin (1:250) and DAPI (1:10,000) for 30min, washed 3X with PBS, and mounted on slides with Fluoromount-G polymerizing medium (Thermo Fisher).

Images for initial experiments (WT, d16, and p95 cell lines) were acquired with a widefield Leica DM300 upright microscope with a DFC345X camera and an HCX PL APO 1003 NA 1.4 objective. Images of at least 75 FOVs per condition were acquired over 3 independent experiments, and Matrix disruption measurements were quantified with FIJI by thresholding matrix disruption area of fluorescent gelatin and dividing by the number of nuclei per field as previously described (PMID 28605388 ). To minimize variation seen across experiments, values were normalized to the WT average of each experiment, and matrix disruption/nuclei were compared as log2-fold change over WT values.

Images for experiments with shRNA knockdowns were acquired with an Axio Imager.Z2 widefield microscope (ZEISS Microscopy, Germany) at 40x magnification, to enable more efficient acquisition of tiled images. Tiled images of at least 2 coverslips per condition for each of 3 independent experiments were analyzed, and total matrix disruption area/nuclei for each coverslip was normalized to the average of WT and compared at log2-fold change over WT average values.

### Cell transplantation-based in vivo assays

#### Orthotopic transplant of cell lines and lung metastasis quantification

Single cell solutions of HCR cell lines were prepared at a concentration of 2 million cells per 50uL of 1:1 Growth Factor-reduced Matrigel (Corning 356231):USP Saline solution. 2 million cells were transplanted bilaterally into the 4^th^ mammary glands of NOD/SCID mice (Jackson Laboratories, NOD.Cg-Prkdc^scid^/J RRID:IMSR_JAX:001303) and monitored by caliper 2x a week until total mouse tumor burden >1600 mm^3 was reached. At sacrifice, lungs and tumors were removed and fixed in 10% neutral buffered formalin (MilliporeSigma HT501128) for at least 24 hours before further processing. To quantify lung metastasis, the protocol for FUnGI lung clarification and imaging was adapted from Rios *et al.* (55) Briefly, FUnGI solution was prepared ahead of time and stored at 4 C. Lungs were removed from fixative solution and washed for at least 30 minutes in 0.5% Tween 20 at room temperature. Lungs were transferred to 5mL Eppendorf tube with 2-3mL of FUnGI solution and placed on a rocker for 24 hours.

Next, lungs were mounted between two glass Micro Slides Superfrost Plus (VWR 48311-703) and imaged with z-dimensions at 10x magnification on the Zeiss 880 confocal microscope. To quantify metastases, maximum intensity projections were created from stitched raw images, underwent channel subtraction to reduce background, then manual annotating in Imaris imaging software. Quantification for all endpoint tumor growth metrics and metastasis was executed in GraphPad Prism.

#### Live imaging of Lifeact-transfected cells

##### Lentiviral packaging and infection

Lentiviral transfer plasmid (pLenti Lifeact-mRuby2, Addgene 84384; or pLenti Lifeact-EGFP, Addgene 84383) and plasmids encoding VSV-G, Gag-Pol, Rev, and Tat genes were transiently transfected into HEK293T cells (ATCC, CRL-11268) with Lipofectamine™ 2000 (Thermo Fisher Scientific). After incubation at 37C for 72h for lentiviral packaging, virus-containing supernatant was collected. Cell lines were infected with virus in media with 5ug/ml of polybrene for 48h, after which virus containing media was removed and cells washed with PBS. LifeAct-infected cell lines were then trypsinized and sorted by FACS to purify fluorescent transfected cells for further propagation in culture for live imaging experiments.

##### Live-cell imaging

Cells were plated on Cellvis 4-chamber 3.5cm glass-bottom dishes (Thermo Fisher Scientific NC0832919) that were pre-treated with 0.1%(w/v in H20) Type B bovine gelatin (Sigma-Aldrich) for 20 minutes, 0.5% glutaraldehyde for 40 minutes, and then rinsed 3x with PBS, and subsequently seeded with 20,000 cells/chamber which were allowed to adhere and grow for 24 hours prior to start of imaging.

Imaging was conducted in phenol-red free DMEM media (Gibco) with 10% FBS (Gemini Bio) and 1% penicillin/streptomycin (Gibco) using an inverted confocal Zeiss LSM 980 microscope with a 37C humidified and temperature-controlled chamber. Z-stacks of 5-6 microns (step size 1um) were acquired every 30-60 min for 10-18 hours at 20x magnification.

##### Quantification and analysis

Timelapse movies of at least 20 FOVs total across at least 2 independent experiments were analyzed, and average number of Invadopodia rosette superstructures per cell at each timepoint were manually annotated in FIJI. Rosette number seen across the total timelapse, normalized to number of nuclei, were totaled for each FOV and compared across conditions.

## Acknowledgements

We acknowledge the members of the Snyder lab, Bravo-Cordero lab, and the Microscopy and Advanced Bioimaging Core at Mount Sinai. JJBC was supported by R01CA244780 (NIH/NCI), NCI R03 (CA270679), R61CA278402 (NIH/NCI), BC241026 (BCRP DOD), the Irma T. Hirschl Trust and the Emerging Leader Award from the Mark Foundation and the Tisch Cancer Institute National Institutes of Health (NIH) Cancer Center grant (P30-CA196521). EB received support from an NIH T32 CA078207 Training Program in Cancer Biology. JCS and HKL acknowledge support from R01CA255372 (NIH/NCI) and BC201085 (BCRP DOD). JDF acknowledges support from NIH T32 5T32HD040372-25 Training Program in Developmental and Stem Cell Biology. SMK acknowledges support from the American Cancer Society Research Scholar award (RSG-23-1039115-01-MM). SMK is a stakeholder in NeoZenome Therapeutics Inc.

